# Orthopoxvirus K3 orthologs show virus- and host-specific inhibition of the antiviral protein kinase PKR

**DOI:** 10.1101/2020.02.20.958645

**Authors:** Chorong Park, Chen Peng, M. Julhasur Rahman, Sherry L. Haller, Loubna Tazi, Greg Brennan, Stefan Rothenburg

**Affiliations:** School of Medicine, University of California Davis, Department of Medial Microbiology and Immunology, Davis, CA 95616, USA; National Institute of Allergy and Infectious Diseases, National Institutes of Health, Laboratory of Viral Diseases, Bethesda, MD 20892, USA; University of Texas Medical Branch at Galveston, Department of Microbiology and Immunology, Galveston, TX 77555, USA

**Keywords:** host-virus interactions, poxvirus, orthopoxviruses, vaccinia virus, variola virus, cowpox virus, PKR

## Abstract

The antiviral protein kinase R (PKR) is an important host restriction factor, which poxviruses must overcome to productively infect host cells. To inhibit PKR, many poxviruses encode a pseudosubstrate mimic of the alpha subunit of eukaryotic translation initiation factor 2 (eIF2), designated K3 in vaccinia virus. Although the interaction between PKR and eIF2α is highly conserved, some K3 orthologs were previously shown to inhibit PKR in a species-specific manner. To better define this host range function, we compared the sensitivity of PKR from 17 mammals to inhibition by K3 orthologs from closely related orthopoxviruses. The K3 orthologs showed species-specific inhibition of PKR and exhibited three distinct inhibition profiles. In some cases, PKR from closely related species showed dramatic differences in their sensitivity to K3 orthologs. Vaccinia virus expressing the camelpox virus K3 ortholog replicated more than three orders of magnitude better in human and sheep cells than a virus expressing vaccinia virus K3, but both viruses replicated comparably well in cow cells. Strikingly, in site-directed mutagenesis experiments between the variola virus and camelpox virus K3 orthologs, we found that different amino acid combinations were necessary to mediate improved or diminished inhibition of PKR derived from different host species. Because there is likely a limited number of possible variations in PKR that affect K3-interactions but still maintain PKR/eIF2α interactions, by chance PKR from some potential new hosts may be susceptible to K3-mediated inhibition from a virus it has never previously encountered. We conclude that neither the sensitivity of host proteins to virus inhibition nor the effectiveness of viral immune antagonists can be inferred from their phylogenetic relatedness but must be experimentally determined.

**Authors summary:** Most virus families are composed of large numbers of virus species. However, in general, only a few prototypic viruses are experimentally studied in-depth, and it is often assumed that the obtained results are representative of other viruses in the same family. In order to test this assumption, we compared the sensitivity of the antiviral protein kinase PKR from various mammals to inhibition by multiple orthologs of K3, a PKR inhibitor expressed by several closely related orthopoxviruses. We found strong differences in PKR inhibition by the K3 orthologs, demonstrating that sensitivity to a specific inhibitor was not indicative of broad sensitivity to orthologs of these inhibitors from closely related viruses. We also show that PKR from even closely related species displayed markedly different sensitivities to these poxvirus inhibitors. Furthermore, we identified amino acid residues in these K3 orthologs that are critical for enhanced or decreased PKR inhibition and found that distinct amino acid combinations affected PKRs from various species differently. Our study shows that even closely related inhibitors of an antiviral protein can vary dramatically in their inhibitory potential, and cautions that results from host-virus interaction studies of a prototypic virus genus member cannot necessarily be extrapolated to other viruses in the same genus without experimental verification.

## Introduction

Poxviruses are a large family of double-stranded (ds) DNA viruses encompassing two subfamilies and 14 currently recognized genera, which include many viruses of both medical and veterinary importance. Poxviruses, as a group, infect a large range of animals; however, the host range of individual virus species can vary dramatically [1]. For example, members of the well-studied orthopoxvirus genus have one of the broadest host ranges of all poxviruses with vaccinia viruses (VACV), cowpox viruses (CPXV) and monkeypox viruses (MPXV) infecting a wide range of mammals. Yet the genus also includes viruses that only infect a single species such as variola virus (VARV), the causative agent of smallpox in humans, and ectromelia virus (ECTV), which only infects mice. There are also orthopoxviruses with a narrow host range such as camelpox virus (CMLV), which primarily infects camels but can also cause infections in humans [2, 3]. This diversity in poxvirus host range is believed to be mediated, at least in part, by a set of host range genes, which have largely evolved to counteract the host antiviral response [1].

One important host antiviral protein that is targeted by poxvirus host range factors is protein kinase R (PKR), which exerts its antiviral effect against many virus families. PKR is ubiquitously expressed in host cells in an inactive monomeric form. In the presence of dsRNA, produced during the replication cycle of most viruses, PKR is activated through a process of dimerization and autophosphorylation. Activated PKR phosphorylates the alpha subunit of eukaryotic translation initiation factor 2 (eIF2), which ultimately prevents delivery of methionyl-tRNA to the ribosome and blocks cap-dependent translation [4-6]. In response to this potent antiviral activity, many viruses have evolved at least one PKR antagonist [7]. These conflicting host and viral proteins exert strong selective pressure on each other, driving rapid, successive changes to the coding sequence of each protein to either disrupt or re-engage the interaction. Signatures of positive selection resulting from this evolutionary “arms race” have been identified across the PKR coding sequence, and some of these positively selected residues have been shown to be determinants of differential susceptibility to both poxvirus and herpesvirus PKR antagonists [8-10].

Many poxviruses express two PKR antagonists, designated E3 and K3 in VACV. E3, the gene product of E3L, contains a N-terminal Z-nucleic acid binding domain and a C-terminal dsRNA binding domain. E3 inhibits PKR by binding to dsRNA and preventing PKR homodimerization [11, 12]. K3, the gene product of K3L, is a structural homolog of eIF2α and acts as a pseudosubstrate inhibitor by binding to activated PKR, thus precluding phosphorylation of eIF2α [13, 14]. Both E3 and K3 have been identified as host range factors, because deletion of either E3L or K3L led to impaired VACV replication in cell lines derived from different species [13, 15]. While the mere presence or absence of host range genes likely contributes to the host range of poxviruses, many orthopoxviruses share a similar profile of host range genes, despite having different host tropisms [16]. This indicates that variations in these host range genes may influence host range by altering interactions with host molecules.

Our recent work has supported this hypothesis in the leporipoxvirus and capripoxvirus genera. We have demonstrated that capripoxvirus K3 orthologs inhibit PKR in a species-specific manner, strongly inhibiting sheep, goat, and human PKR, while only weakly inhibiting cow and mouse PKR [17]. Similarly, M156, the K3 ortholog from the rabbit- and hare-specific myxoma virus (MYXV), inhibited European rabbit PKR but not PKR derived from seven other mammals. Moreover, this work demonstrated that changes in viral antagonists can be rapidly fixed in the population, as demonstrated by the discovery that M156 evolved a loss of function mutation that might contribute to the attenuation of the virus in European rabbits [18].

Most research on orthopoxviruses has focused on the prototypic poxvirus VACV as a model for poxvirus virology and host-virus interactions. However, very few studies have systematically investigated the activity of VACV orthologs in other orthopoxviruses. It is therefore unclear whether the activity of VACV proteins is representative of their orthologs in other orthopoxvirus. Orthopoxviruses such as CPXV and VARV represent substantial public health or bioterror threats, and human infections with orthopoxviruses are predicted to become more common as immunity due to the cessation of the smallpox vaccination decreases in the population [19]. Therefore, it is critically important to identify orthopoxvirus determinants of host range and to identify viruses most likely to infect new species.

In this study, we compared inhibition of PKR from a panel of mammalian species by K3 orthologs from VACV, VARV, CMLV and CPXV-rat09 to determine whether the ability of VACV K3 to inhibit PKR from a given host was predictive of the ability for other orthopoxvirus K3 orthologs to inhibit this host antiviral protein. The results presented here show that the tested K3 orthologs exhibited distinct inhibition profiles and also that the phylogenetic relatedness of PKR was not a good predictor for how susceptible they were to inhibition by K3 orthologs. Mapping K3 residues that governed the species-specificity of these interactions revealed that multiple amino acid substitutions were necessary to convert a weak K3 ortholog into a better inhibitor of most tested PKRs. Furthermore, different amino acid combinations were needed to most effectively inhibit PKR from different species.

## Results

### Species-specific inhibition of mammalian PKR by vaccinia virus K3

Previous yeast-based and transfection assays indicated differential inhibition of PKR from several vertebrate species by VACV K3 [8, 9]. In order to get a more complete picture about the breadth of species-specific PKR inhibition, we analyzed the sensitivity of PKRs from 17 mammals to inhibition by VACV K3 in an established luciferase-based reporter (LBR) assay (Fig. 1A) [9]. In agreement with previous findings, human, and sheep PKRs were largely resistant, whereas mouse and cow PKRs were sensitive to K3 inhibition in this assay [17]. Among the primate PKRs tested, tamarin PKR was most strongly inhibited, consistent with previous yeast assays [8]. Because vaccinia virus is thought to have originated from a rodent virus, it was surprising that only Syrian hamster and mouse PKRs were sensitive, whereas PKRs derived from other rodents, rats and guinea pigs, were resistant. Among PKRs from ungulates, horse, cow and camel PKR were sensitive, whereas sheep PKR was only weakly sensitive and pig PKR was resistant. The observation of such extensive variability in sensitivity to VACV K3, even among related species, raised the question as to whether other orthopoxvirus K3 orthologs would follow this pattern of PKR inhibition.

**Figure 1.**
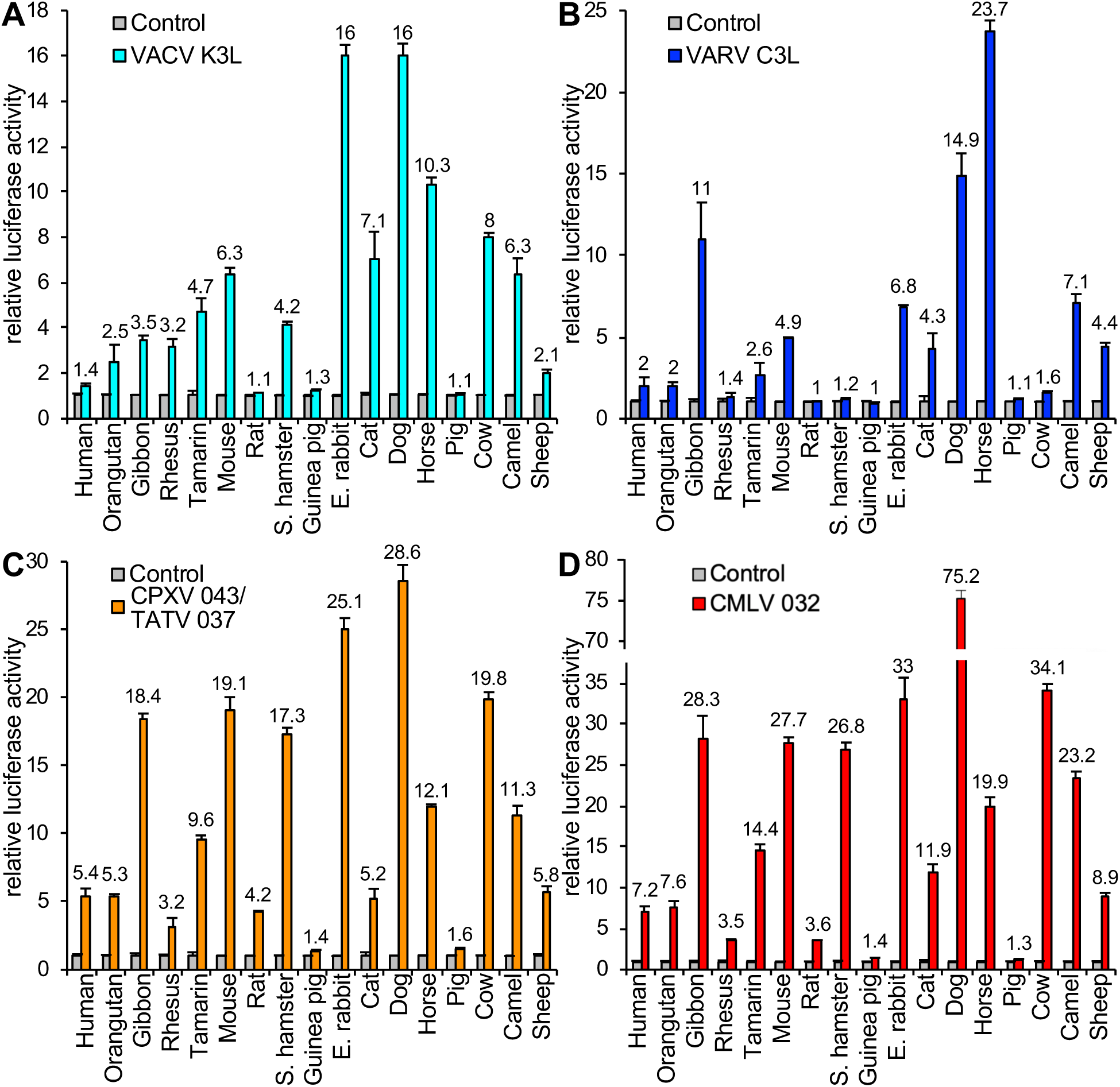
Species-specific inhibition of PKR by orthopoxvirus K3 orthologs. HeLa-PKR^kd^ cells were transfected with expression vectors encoding firefly luciferase (0.05 μg), PKR from the indicated species (0.2 μg) and either 0.2 μg VACV K3L (**A**) or VARV C3L (**B**), CPXV-D 043/TATV 037 (**C**), or CMLV 032 (**D**). Luciferase activities were measured 48 hours after transfection and normalized to PKR-only transfected cells to obtain relative luciferase activities. Error bars represent the standard deviations from three independent transfections. Results shown are representative of at least three independent experiments.

### Inhibition profiles of K3 orthologs from other orthopoxviruses are distinct from VACV K3

To compare the inhibition profiles of VACV K3 orthologs from other orthopoxviruses, we cloned VARV C3L, CMLV (CMS strain) 032, and CPXV 043 from the CPXV-rat09 strain, the latter of which is identical to multiple CPXV isolates from the D clade [20, 21], into the mammalian expression vector pSG5. CPXV-rat09 043 is also identical to protein 037 from taterapox virus (TATV) and will henceforth be referred to as CPXV-D 043/TATV 037. Relative to VACV K3, CPXV-D 043/TATV 037 has 11 amino acid (aa) differences (87.5 % identity), CMLV 032 has 12 aa differences (86.4 % identity), and VARV C3 has 17 aa differences (80.7 % identity) (Fig. 2A, B). Transient transfection of HeLa-PKR^kd^ cells with the K3L orthologs revealed comparable expression by Western blot (Fig. 2C). These three K3 orthologs showed different inhibition profiles by the LBR assay, which were distinct from each other and from the VACV K3 inhibition pattern (Fig. 1 and Fig. 3). Compared to VACV K3, VARV C3 showed slightly better inhibition of human and horse PKR, and considerably better inhibition (> 2-fold) of gibbon and sheep PKR. In contrast, VARV C3 inhibited PKR from eight other species less well than VACV K3. CPXV-D 043/TATV 037 and CMLV 032 showed overall similar inhibition profiles to one another, inhibiting PKR from most species better than VACV K3. However, there was essentially no difference in the ability of CPXV-D 043/TATV 037 or CMLV 032 to inhibit rhesus, cat, horse or camel PKR, compared to VACV K3. Pig and guinea pig PKR were largely resistant to inhibition by all K3 orthologs tested.

**Figure 2.**
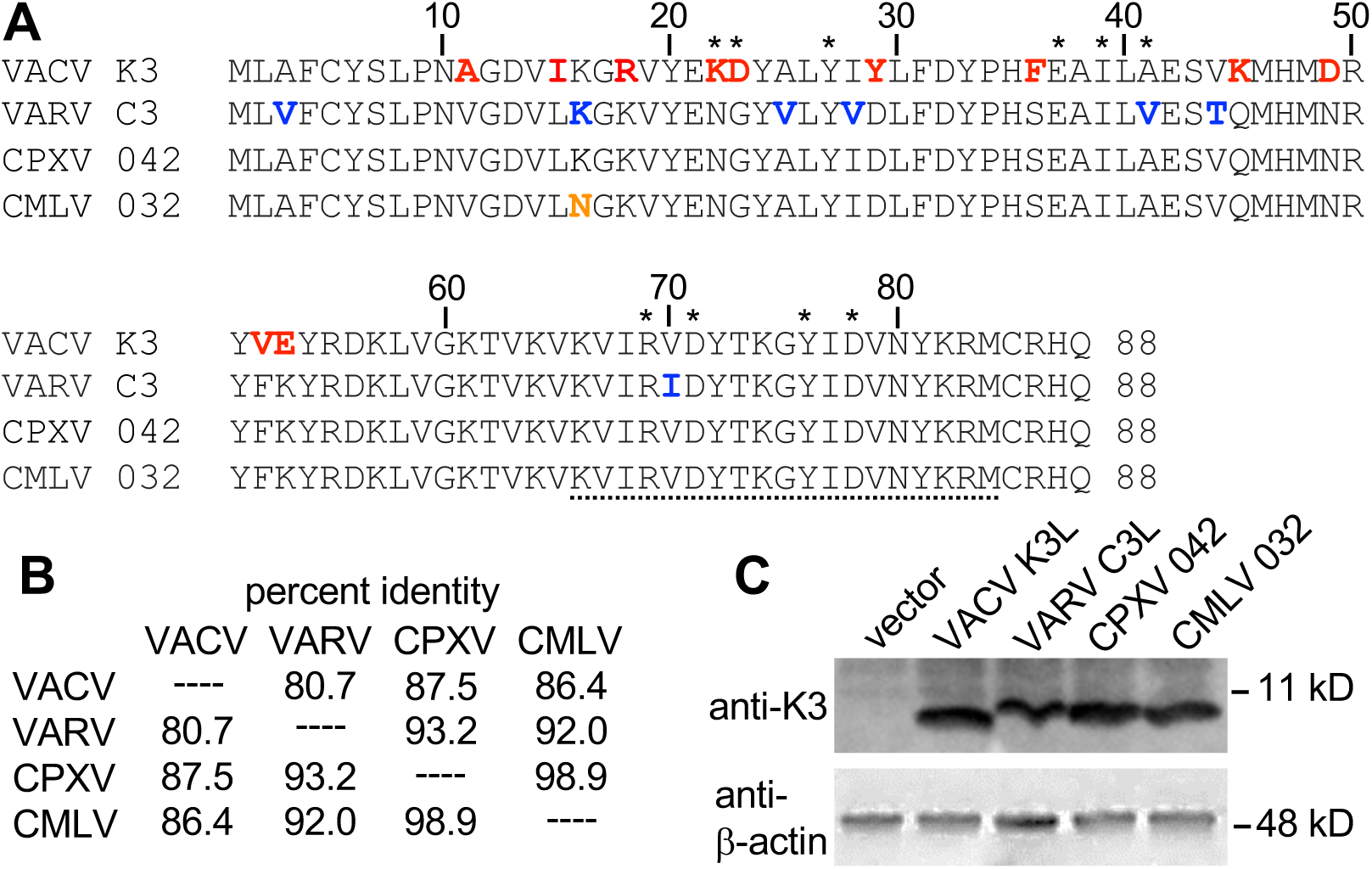
Amino acid differences between orthopoxvirus K3 orthologs. (**A**) Multiple sequence alignment of VACV K3, VARV C3, CPXV-D 043/TATV 037, and CMLV 032. Amino acids that are unique to VACV K3 are shown in red. Residues that differ between VARV C3 and CMLV 032 are highlighted in blue. The residue that is unique to CMLV 032 is shown in orange. Asterisks denote residues of K3 that correspond to the contacts of eIF2α with PKR in a co-crystal structure [51]. The peptide sequence that was used to generate anti-VACV K3 antibodies is underlined. (**B**) Amino acid identities between tested K3 orthologs used in this study. Percent identities were calculated from a multiple sequence alignment using MegAlign. (**C**) Expression of K3 orthologs in transfected cells. HeLa-PKR^kd^ cells were transfected with the indicated K3L orthologs, and cell lysates were collected 48 hours later. Total proteins from these samples were separated on 12% SDS-PAGE gels and analyzed by immunoblot analysis with the indicated anti-K3 and anti-β-actin antibodies.

**Figure 3.**
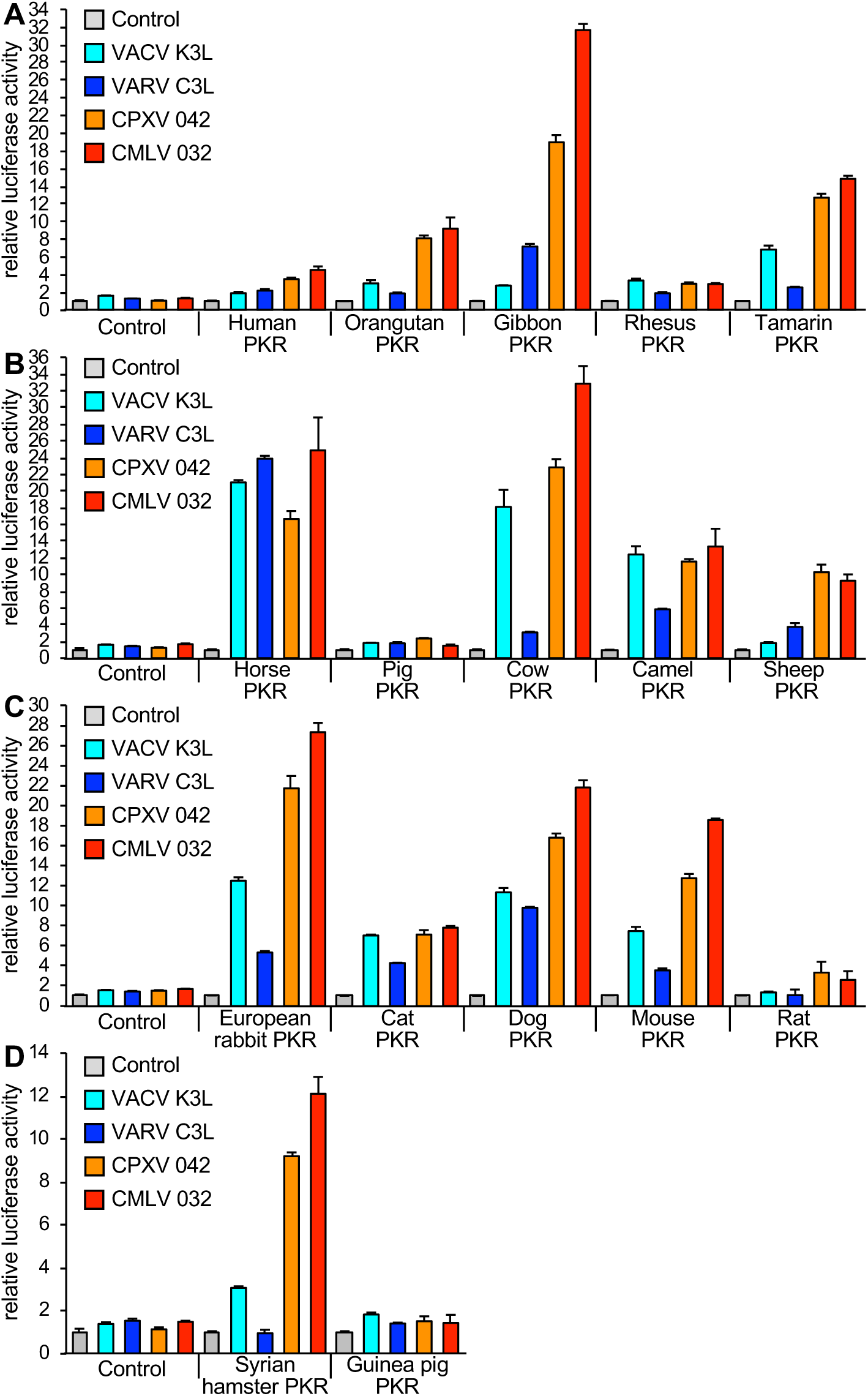
Orthopoxvirus K3 orthologs show differential PKR inhibition. HeLa-PKR^kd^ cells were transfected with expression vectors encoding firefly luciferase (0.05 μg), with either VACV K3L, VARV C3L, CPXV-D 043/TATV 037, or CMLV 032 (0.4 μg), and PKR (0.2 μg) from the indicated species in four groups (**A-D**). Luciferase activities were measured as described in Fig. 1. Error bars represent the standard deviations from three independent transfections. Results shown are representative of at least three independent experiments.

To visualize if the phylogenetic relatedness of a given PKR may be predictive of susceptibility to these K3 orthologs, we generated a phylogenetic tree using PKR sequences derived from 40 mammalian species. We included PKRs from several species that were not tested in the LBR assays to achieve better statistical support for the branches. We then mapped the relative sensitivities of the tested PKRs to each K3 ortholog obtained from multiple LBR assays using a susceptibility scale ranging from weakly sensitive to highly sensitive on a scale from 1 to 6, respectively (Fig. 4). By this analysis there was no discernible pattern to predict PKR sensitivity by phylogenetic relatedness.

**Figure 4.**
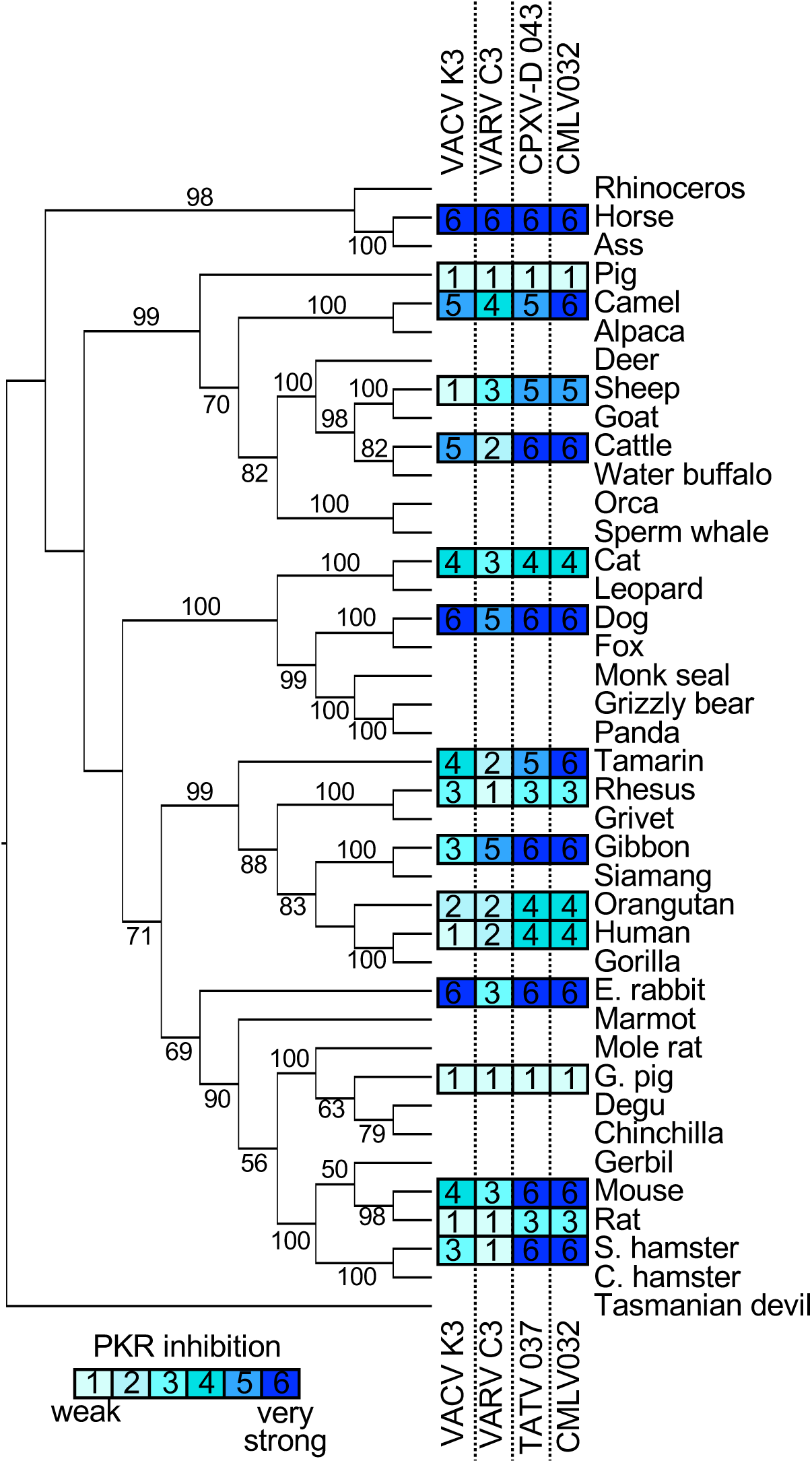
Sensitivities of tested PKRs projected on a PKR phylogenetic tree. A maximum-likelihood phylogenetic tree of PKR from selected mammalian species including the ones tested in this study was generated. The tree was rooted to Tasmanian devil PKR. Bootstrap values ≥ 50 are indicated on the branches. Relative sensitivities of PKRs to VACV K3, VARV C3, CPXV-D 043/TATV 037, or CMLV 032 obtained from multiple experiments with PKR:K3 ortholog ratios of 1:1 and 1:2 are shown on a scale from 1 to 6: 1 = no or very weak inhibition (≤ 2 fold); 2 = weak inhibition (2 to 3-fold); 3 = intermediate low inhibition (3 to 5-fold); 4 = intermediate high inhibition (5 to 7 fold); 5 = high inhibition (7 to 10-fold); 6 = very high inhibition (≥ 10-fold).

### Sensitivity of PKR to K3-mediated inhibition correlates with host range function during infection

In order to analyze whether the potency of PKR inhibition by VACV and CMLV K3 orthologs correlates with the ability of these proteins to support productive infection, we cloned VACV K3L and CMLV 032 into the E3L locus of the highly attenuated VACV strain VC-R4, which lacks both K3L and E3L and can only replicate in the absence of PKR or in cells that express exogenous PKR inhibitors [22]. Using these viruses, we compared plaque formation and virus replication relative to wild type vaccinia virus (VC-2, Copenhagen strain) in cells of human (HeLa), gibbon (Tert-GF), sheep (OA1) or cow (BT) origin. In HeLa cells that have PKR knocked out (HeLa-PKR^ko^), all four viruses formed comparably sized plaques (Fig. 5A). In contrast, infected wild type HeLa or Tert-GF cells only developed plaques when they were infected with VC-2 or VC-R4+CMLV 032. These viruses also caused large plaque formation in OA1 cells, whereas infection with VC-R4+VACV K3L led to the formation of small plaques. In BT cells, all viruses, except for VC-R4, formed comparably sized plaques.

**Figure 5.**
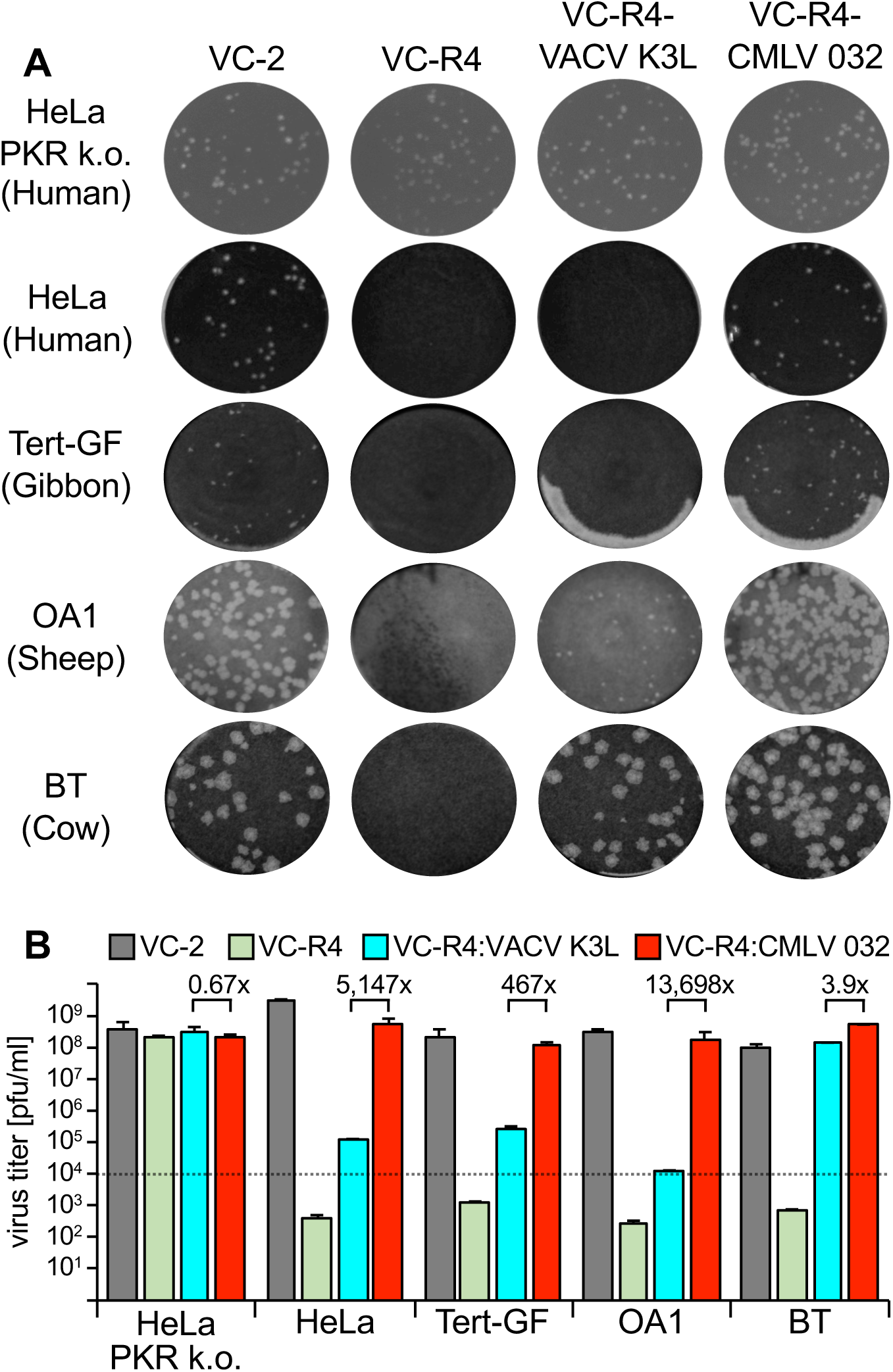
CMLV 032 facilitates strong VACV replication. HeLa-PKR^ko^, HeLa, Tert-GF, OA1 and BT cells were infected with VC-2 (containing E3L and K3L), VC-R4 (missing E3L and K3L), VC-R4:VACV_K3L, and VC-R4:CMLV_032. (**A**) Cells were infected with each virus for 1 hour and the overlayed with DMEM + 2% carboxymethylcellulose to allow plaque formation. Cells were stained with 0.1% w/v crystal violet to visualize plaque size two days post-infection. (**B**) Cells were infected with the indicated viruses at MOI = 0.01. 48 hours after infection, viruses in cell lysates were tittered on RK13+E3L+K3L cells. Error bars represent the standard deviations from two independent infections. Fold differences in virus titers obtained with VC-R4:VACV_K3L, and VC-R4:CMLV_032 are shown.

Multiple cycle replication assays also correlated with the results of the plaque assays, with VC-2, VC-R4+CMLV 032, and VC-R4+VACV K3L replicating to comparable titers in HeLa PKR^ko^ cells and BT cells. In Contrast, VC-2 and VC-R4+CMLV 032 were the only viruses that replicated well in HeLa, Tert-GF, and OA1 cells. Viruses containing VACV-K3L replicated to slightly higher titers than input (∼10-fold increase) in HeLa and Tert-GF cells indicating that the weak inhibition of PKR allowed limited virus replication.

### Identification of amino acid residues that confer differential PKR inhibition between VARV C3L and CMLV 032

VARV C3L and CMLV 032 differ by seven aa residues (Fig. 2). In order to identify which residues are responsible for the virus-specific differences in PKR inhibition, we first exchanged single aa residues in VARV C3L with the ones found in CMLV 032 and performed the LBR assay with human, tamarin, mouse, dog, cow and sheep PKR. None of these mutants inhibited human PKR better, whereas some mutants showed better inhibition of some other PKRs. Only one VARV C3L mutant (V25A) reached comparable PKR inhibition as CMLV 032, and only for dog PKR. VARV C3L-V25A showed more than 2-fold increased PKR inhibition of tamarin and sheep PKR and more than 50% higher inhibition of cow and mouse PKR. VARV C3L-I70V showed more than 2-fold increased PKR inhibition of tamarin PKR and more than 50% higher inhibition of mouse and dog PKR. VARV C3L-V3A displayed more than 50% higher inhibition of dog and sheep PKR, whereas VARV C3L-T44V only showed higher inhibition of mouse PKR (more than 2-fold).

These experiments confirmed that a single aa residue was not responsible for the differential PKR inhibition found for most of the tested PKRs. Therefore, we generated mutants containing multiple aa changes, designated as M2-M7 (Fig. 7A). The majority of these residues localized to the suspected PKR-K3 interacting face when we mapped these positions on the existing VACV K3 structure (blue and purple, Fig. 7B). We expressed each mutant VARV C3L in transfected HeLa-PKR^ko^ cells and confirmed expression by Western blot analysis. VARV C3L and M1 were detected slightly less well, which might be explained by the presence of isoleucine at position 70, potentially altering the epitope detected by the primary antibodies. Mutation of this residue to valine (M2 and up), which is included in the native epitope, resulted in detection comparable to CMLV 032. Remarkably, all tested PKRs showed different sensitivity patterns (Fig. 7D). For human PKR, a combination of four mutations (M4) resulted in inhibition comparable to CMLV 032, while mutations in M2 and M3 had no effect. Inhibition of tamarin PKR increased more gradually, with M4 being as effective as CMLV 032. Inhibition of mouse PKR increased sharply with the inclusion of T44V in M6. Inhibition of cow PKR showed the most gradual, stepwise increase from M1 to M7.

**Figure 6.**
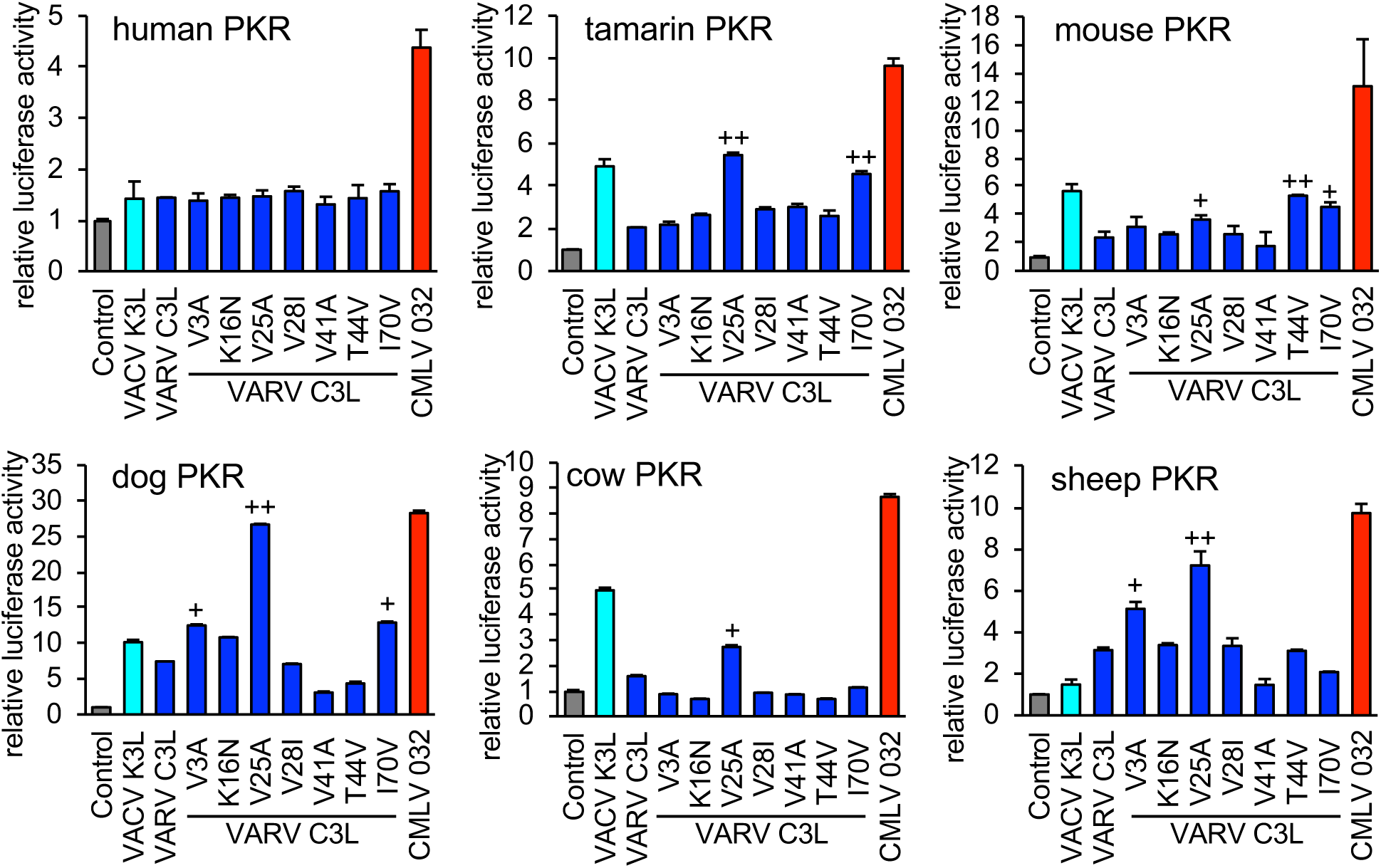
Identification of amino acid residues in VARV C3 that confer differential PKR inhibition. HeLa-PKR^kd^ cells were transfected with expression vectors encoding firefly luciferase (0.05 μg), with either VACV K3L, VARV C3L, the indicated mutants of VARV C3L, or CMLV 032 (0.4 μg), and PKR from the indicated species (0.2 μg). Luciferase activities were measured 48 hours after transfection and normalized to PKR-only transfected cells to obtain relative luciferase activities. Error bars represent the standard deviations from three independent transfections. Results shown are representative of three independent experiments. 1.5-fold and 2-fold (+) and ≥2 fold (++) higher inhibition of the mutant than VARV C3L wild type are indicated.

**Figure 7.**
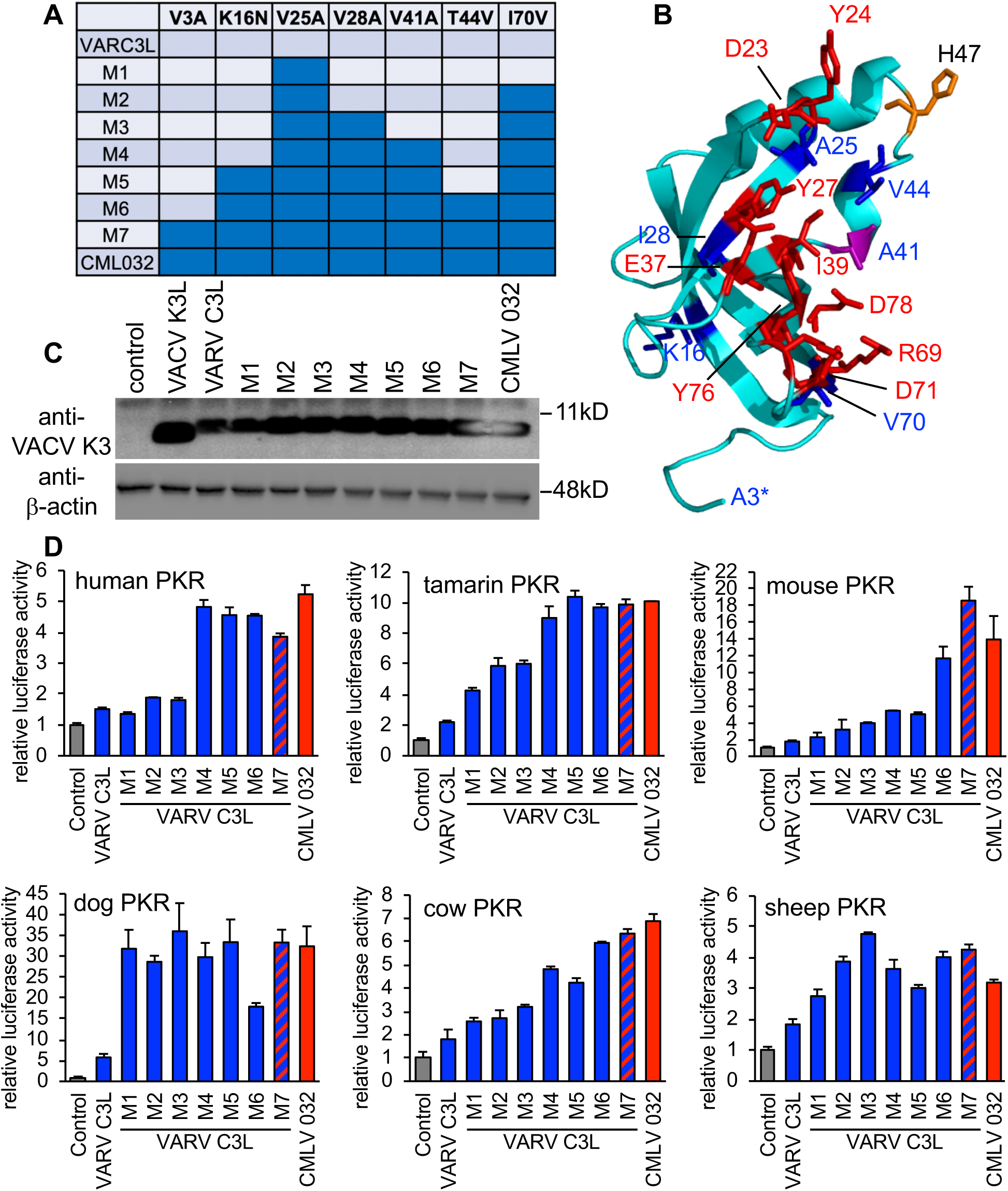
Distinct amino acids combinations are responsible for the higher inhibition of CMLV 032 for PKRs from different species. (**A**) Seven different mutants of VARV C3L were generated that contain successive combinations of amino acid substitutions that are found in CMLV 032. (**B**) Crystal structure of VACV K3 [14]. Residues that correspond to amino acids of eIF2α that contact human PKR in a co-crystal structure [51] are shown in purple (residue A41) or red (other residues). Residues that differ between VARV C3 and CMLV 032 are shown in purple (residue A41) or blue (other residues). Residue H47, which has been previously implicated in PKR binding [52], is shown in orange. Residues in eIF2α corresponding to residues 1-3 (asterisk) 44-52 in K3 were not resolved in the co-crystal structure. (**C**) Expression of VARV C3 mutants, CMLV 032 and VACV K3 in transfected HeLa-PKR^kd^ cells. Cell lysates from transfected cells were separated on 12% SDS-PAGE gels and analyzed by immunoblot analysis with anti-K3 and anti-β-actin antibodies. (**D**) HeLa-PKR^kd^ cells were transfected with expression vectors encoding firefly luciferase (0.05 μg), with VARV C3L, the indicated mutants of VARV C3L, or CMLV 032 (0.4 μg), and PKR (0.2 μg) from the indicated species. Luciferase activities were measured 48 hours after transfection and normalized to PKR-only transfected cells to obtain relative luciferase activities. Error bars represent the standard deviations from three independent transfections. Results shown are representative of two independent experiments.

**Figure 8.**
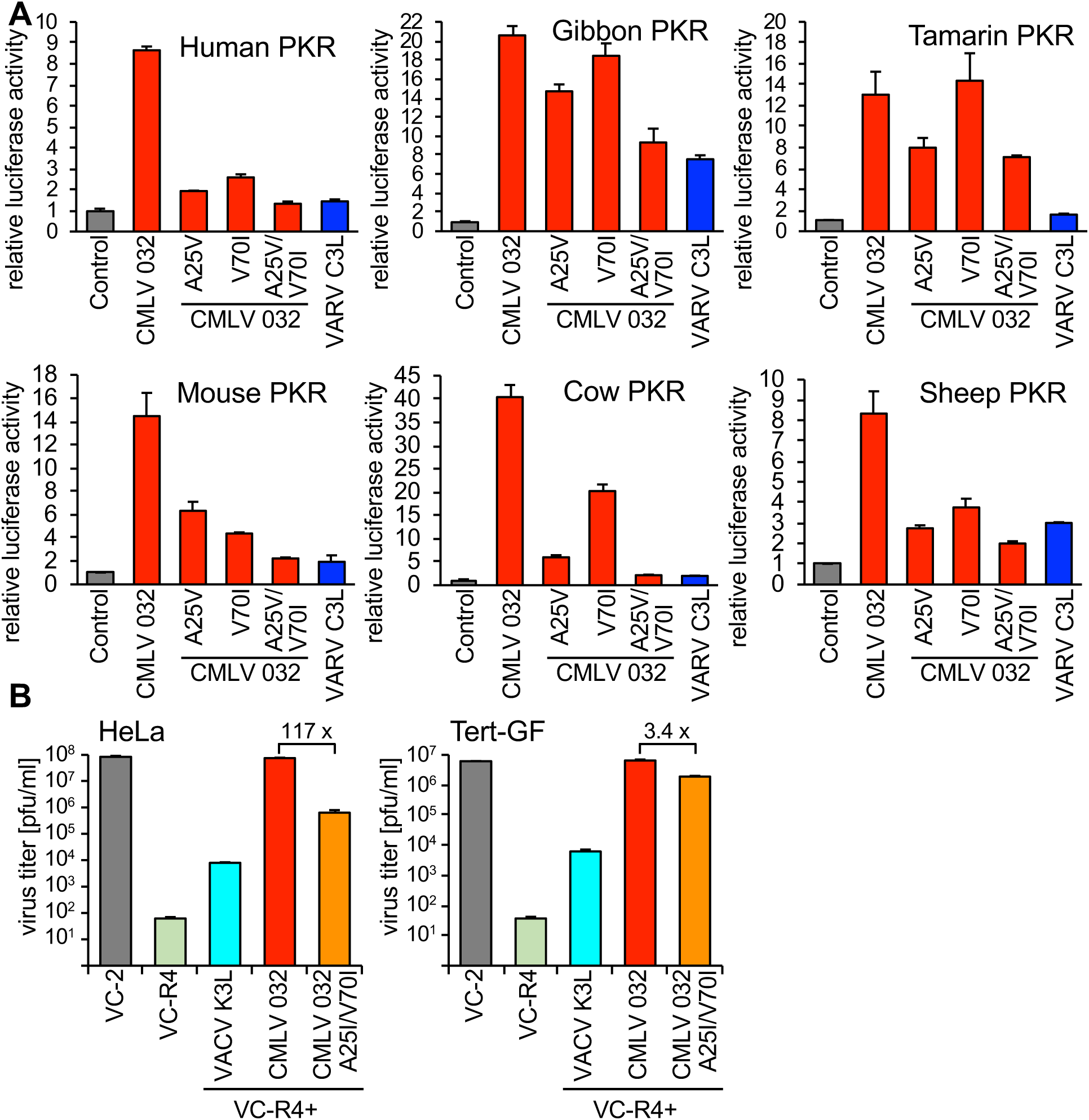
Residues A25 and V70 in CMLV 032 are critical for strong PKR inhibition. (**A**) HeLa-PKR^kd^ cells were transfected with expression vectors encoding firefly luciferase (0.05 μg), with CMLV 032, the indicated mutants of CMLV 032, or VARV C3L (0.4 μg), and PKR from the indicated species (0.2 μg). Luciferase activities were measured 48 hours after transfection and normalized to cells transfected only with PKR to obtain relative luciferase activities. Error bars represent the standard deviations from three independent transfections. Results shown are representative of two independent experiments. (**B**) HeLa or gibbon (Tert-GF) cells were infected with VC-2, VC-R4 or its derivatives containing either VACV K3, CMLV 032 or CMLV 032A25I/V70I at a MOI of 0.01. After 48 hours viruses were harvested and tittered on RK13+E3L+K3L cells. Error bars represent the standard deviations of two independent experiments. Fold differences in titer between VC-R4+CMLV 032 and VC-R4+CMLV 032-A25I/V70I are shown.

### Alanine 25 and isoleucine 70 in CMLV 032 are important for strong PKR inhibition

Because swapping residues 25 and 70 in VARV C3L with the ones in CMLV 032 resulted in better inhibition of PKRs from many species, we replaced the corresponding residues in CMLV 032 with the VARV C3L residues either alone or in combination. Single mutations led to strongly decreased inhibition of human, mouse, cow and sheep PKR. For these species’ PKR, the double mutant CMLV 032-A25V/V70I, showed PKR inhibition comparable to VARV C3L, indicating additive effects of both mutations. Single CMLV 032 mutants changed the inhibition of gibbon PKR only weakly, and replacement of both A25V and V70I resulted in inhibition comparable to VARV C3L. Tamarin PKR was the only PKR tested for which the double-mutant displayed substantially stronger PKR inhibition than VARV C3L. We next compared replication of VACV containing CMLV 032-A25V/I70L with a virus expressing either CMLV 032 or VACV K3L. Correlating with the LBR assays, VACV-CMLV 032-A25V/I70L showed only modestly lower (3.4-fold) replication than VACV-CMLV 032 in tert-GF, whereas 117-fold lower replication was observed in HeLa cells. Notably, VACV-CMLV 032-A25V/I70L showed higher replication than VACV-K3L in HeLa cells.

## Discussion

Most research on poxviruses has focused on VACV, which is often called the prototypic poxvirus. VACV K3 has been shown to inhibit PKR from several mammals in a species-specific manner, as defined in both LBR assays and yeast-based studies [8, 9, 17]. The data presented here support and extend this species-specific sensitivity of PKR to VACV K3L and add to a growing body of evidence that virus-specific differences play an important role in viral host range, as do species-specific differences in host genes [23, 24]. Consistent with this concept, in this study we show that the VACV K3 orthologs from four different orthopoxviruses displayed dramatic differences in their ability to inhibit PKR from different species, including different sensitivities in closely related species such as mice and rats. Furthermore, VACV, VARV and CMLV/CPXV-D 043/TATV 037 orthologs showed three distinct inhibition profiles when tested against 17 mammalian PKR orthologs. The extent of this phenomenon across orthopoxvirus K3L orthologs uncovered here was unanticipated.

The three distinct patterns of PKR inhibition mediated by K3 orthologs that we identified in this study do not appear to be predictable by either phylogenetic distance between different PKRs, or by whether the PKR is derived from a natural or alternative host species. For example, among the primate PKRs tested, VACV K3L only inhibited tamarin PKR efficiently, although VACV can infect many primates [25]. However, VARV K3 ortholog C3L only inhibited gibbon PKR efficiently. Surprisingly, despite VARV being a human-specific virus, C3L did not efficiently inhibit human PKR. In contrast, although CMLV does not efficiently infect primate species, CMLV 032 inhibited all primate PKRs except rhesus PKR. The inhibition of PKRs by VACV K3 and CMLV 032 in the LBR assays correlated with the ability of those inhibitors to support virus replication *in vitro*. A VACV expressing only K3L replicated three to four orders of magnitude better in bovine cells than in human, gibbon, or sheep cells. In contrast, the same virus expressing the CMLV K3 ortholog 032 replicated as efficiently as wild type VACV in all of these cell types. This also demonstrates that one efficient K3 ortholog can substitute for both VACV E3 and K3, in at least some cell lines.

The strong activity of CMLV K3 against PKR from its natural host (camel) is in line with previous studies that showed that myxoma virus, sheepox virus and goatpox virus inhibited PKR from their naturals hosts with high efficiency [17, 18]. In contrast, we have shown here and in a previous study that K3 orthologs from two other poxviruses, lumpy skin disease virus and VARV, only weakly inhibited PKR from their natural hosts, cattle and humans, respectively [17]. Because CPXVs possess very broad host ranges it is difficult to make conclusions about their natural hosts [25]. As of now, 26 CPXV isolates possess K3 orthologs that are identical to CPXV-D 043/TATV 037 on the protein level. These viruses were mostly isolated from humans (38%), rats (31%) and cats (23%) [20, 26, 27], and PKR derived from these hosts were intermediately sensitive to this K3 ortholog. The host range of TATV, which was isolated from an healthy gerbil (*Tatera kempi*), is unknown [28].

It is not clear, why some poxvirus K3 orthologs are efficient inhibitors of their host’s PKRs, whereas others are not. A possible explanation is that different poxviruses may produce different amounts of dsRNA during infection. The formation of dsRNA is an important danger signal detected by multiple innate immune system sensors, and perhaps as a consequence, the amount of dsRNA produced during infection varies even between closely related orthopoxviruses. For example, vaccinia virus produces high amounts of dsRNA during infection, while monkeypox virus (MPXV) and ectromelia virus (ECTV) produce lower amounts of dsRNA [29, 30]. In agreement with this observation, total RNA produced during MPXV infection was a less potent activator of human PKR than total RNA produced during VACV infection [29]. Strikingly, MPXV and ECTV contain inactivating mutations in their K3L orthologs [16]. Thus, while K3 may be a necessary PKR antagonist in viruses that produce large amounts of dsRNA, possibly releasing some E3 to antagonize other dsRNA-activated pathways, poxviruses that naturally produce less dsRNA might not require redundant PKR inhibitors. It will be interesting to determine whether this correlation between reduced dsRNA production and K3 activity holds across multiple poxviruses.

In many cases, K3 orthologs inhibited PKR from species that are not the natural hosts of the respective poxvirus, e.g. CMLV 032 strongly inhibited gibbon, dog and cow PKR, even though these species are likely never infected by CMLV. There is evidence of substantial positive selection in PKR throughout the mammalian phylogeny, likely driven by a diverse set of viral PKR inhibitors [8, 9]. These positively selected residues cluster around the eIF2α binding region of PKR. Because the PKR/eIF2α interaction must be preserved, there are likely a limited number of evolutionary options available to disrupt this interface between PKR and pseudosubstrate inhibitors like K3. Thus, even though CMLV 032 likely evolved to inhibit PKR in the camelid lineage, the evolutionary constraints on PKR increase the likelihood that PKR from other species might by chance be susceptible to this inhibitor. In line with this is a recent report that shows that VACV strains that contain different poxvirus K3 orthologs as their only PKR inhibitors exhibited different replication profiles in cell lines from different species, with some strains showing broader host ranges, whereas others showed narrower host ranges [31]. In a broader context, these chance susceptibilities of host antiviral factors to viral inhibitors that a host has otherwise never been exposed to, might play an important role in cross-species transmission of viruses [24].

Although VARV is a human-specific pathogen, this study and previous yeast-based assays demonstrated that the VARV K3L ortholog does not inhibit human PKR efficiently [8, 9]. However, mutation of some of the positively selected residues in human PKR to residues found in other species differentially affected the sensitivity to VACV K3 and VARV C3, further illustrating that these differences are mediated by only a few residues in either the host or viral protein [9]. The close relatedness between VARV C3 and CMLV 032 allowed the identification of aa residues that are responsible for the differential inhibition of PKR. In 5 of the 6 tested PKR mutants, single aa substitutions in VARV C3 led to little or no change in activity. Distinct combinations of mutations were necessary to make VARV C3 as efficient as CMLV 032 depending on the particular PKR ortholog being tested. For example, only V25A was necessary to make VARV C3 inhibit dog PKR as efficiently as CMLV 032; however, four aa changes were necessary to phenocopy CMLV 032 inhibition of human PKR by VARV C3, and only three aa changes were necessary for this effect against sheep PKR. It is noteworthy that the loss of PKR inhibition by CMLV 032 was more easily achieved by single aa substitutions than an increase in PKR inhibition. The differences in VARV C3L were all acquired after the split from a CMLV/TATV ancestor, and all of these C3L mutations are non-synonymous, which is indicative of positive selection [8]. This might indicate that there was some selective advantage for VARV C3 to become a less efficient inhibitor of human PKR. Alternatively, these mutations might have occurred in an ancestor of VARV as an adaption to a different host, before it became human-specific.

In addition to the work on K3 orthologs, there is evidence that other poxvirus immune antagonists can have varying effects on the host. For example, while the E3 orthologs derived from swinepox virus and myxoma virus rescued E3L-deficient VACV in HeLa cells, the sheeppox virus E3 ortholog was not able to rescue this virus [32]. Along this line, sheeppox virus E3L was not effective in inhibiting sheep PKR in the LBR assay [17]. These findings indicate that not all E3L orthologs are as broadly functional as currently believed. Thus, it is possible that poxviruses whose E3 orthologs inhibit PKR less efficiently might depend more on their K3 orthologs for PKR inhibition than is currently recognized. Similarly, orthologs of the complement-binding factor VCP from VACV, VARV and MPXV were shown to inhibit complement factors in a species-specific manner [33-37]. Another example of variations in poxviral immune antagonist activity is found in the poxvirus C7L family, which confers host range function in different poxviruses [38-40]. C7 orthologs were shown to bind and inhibit host restriction factors SAMD9 and SAMD9L in a species-specific manner [41-43]. Interestingly, SAMD9/SAMD9L inhibition can be mediated by different, unrelated poxvirus proteins such as CPXV CP77 or K1 in some poxviruses. CPXV CP77 was recently shown to bind and inhibit Chinese hamster SAMD9L, which was not inhibited by VACV C7 or K1 [44]. In combination with the data we present in this study, these results emphasize the fact that immunomodulation is an important component of species-specificity. Changes in a small number of residues in viral immunomodulators may confer unexpected sensitivities to antiviral factors from species that have not encountered the virus. This chance sensitivity may simply be due to the relatively small number of residues that appear to govern these host-virus interactions. In some cases, exemplified by SAMD9/SAMD9L inhibition, virus-specific differences in strategies to inhibit a particular host immune response adds a further layer of complexity to the task of predicting virus host range *a priori*. Ultimately, it is likely that a combination of some or all of these interactions is necessary to permit a virus to replicate in a given host [24].

Species-specific interactions of PKR with viral inhibitors have also been observed with cytomegalovirus (CMV) proteins. Human CMV TRS1 only inhibited human PKR but not PKR from old world monkeys, while the TRS1 ortholog from African green monkey (AGM) CMV inhibited AGM but not human PKR. Intriguingly, mutation of a single amino acid in human PKR (position 489), which determines susceptibility to human or AGM TRS1, is also involved in the weak sensitivity of human PKR to VACV K3 inhibition. Taken together, these data highlight that inhibitors from different pathogens can be affected by mutations in individual antiviral proteins [8, 10].

Large DNA viruses commit about half of their genes to counteract and redirect the host immune response. Because many viruses evolve with specific hosts, it can be assumed that a considerable portion of their encoded proteins will show some level of host-specificity, which ultimately contributes to their host range and virulence. Our study shows that orthologs of even closely related viruses can show dramatic differences in their interaction with host innate immune system. Furthermore, the phylogenetic relatedness of PKR was not a good predictor for how well they were inhibited by K3 orthologs. These findings imply that research obtained with one specific virus and one specific host species cannot necessarily be generalized, instead these interactions likely require experimental validation for other viruses and species.

## Materials and methods

### Cell lines

HeLa (human, ATCC #CCL-2), HeLa PKR-knockdown(^kd^) (kindly provided by Dr. Charles Samuel [45]), HeLa PKR-knockout (kindly provided by Dr. Adam Geballe [10]), RK13+E3L+K3L cells (rabbit) [46], tert immortalized gibbon fibroblasts (tert-GF, kindly provided by Dr. Stephanie Child and Dr. Adam Geballe), OA1 (sheep, ATCC #CRL-6538) and BT (cow, ATCC #CRL-1390) were maintained in Dulbecco’s Modified Eagle’s Medium (DMEM) supplemented with 5% fetal bovine serum or 10% horse serum (BT cells) and 100 IU/ml penicillin/streptomycin (Gibco). HeLa PKR^kd^ cells were maintained in medium containing 1 µg/ml puromycin (Sigma). RK13+E3L+K3L cell culture medium contained 500 µg/ml geneticin and 300 µg/ml zeocin (Life Technologies).

### Plasmids

All PKR and viral genes were cloned into the pSG5 expression vector (Stratagene) for transient transfection assay. Cloning of cow and sheep PKR, European rabbit PKR, mouse PKR, rat PKR, Syrian hamster PKR and guinea pig PKR was described previously [18]. Orangutan PKR, gibbon PKR, macaque PKR, and tamarin PKR were kindly provided by Dr. Nels Elde [8]. Synonymous mutations were introduced in the target site for the shRNA in orangutan PKR, gibbon PKR, macaque PKR and tamarin PKR to generate knock-down resistant PKRs. Horse PKR was cloned from efc cells (kindly provided by Dr. Udeni Balasuriva and Dr. Ying Fang). Pig PKR was cloned from PK15 cells (ATCC-CCL-33). Camel PKR was cloned from Dubca cells (ATCC-CRL2276). Cloning of VACV_K3L (YP_232916.1) was previously described [9]. VARV_C3L (DQ441439.1) and CPXV-D 043/TATV 037 (AGY97204.1/YP_717344.1) were cloned into pSG5 from synthesized DNA. CMLV_032 (NP_570422.1) was generated from pSG5-CPXV-D_043/TATV_037 by site-directed mutagenesis. VACV K3L and CMLV 032 constructs were cloned into P837-GOI-mCherry-E3L for recombinant VACV generation using primers containing *SaI*I (forward primer) and *Nhe*I (reverse primer) sites, as described [22]. VARV C3L and CMLV 032 single and overlapping mutations were generated by site-directed mutagenesis using Phusion High-Fidelity DNA polymerase (NEB). All ORFs were validated by Sanger sequencing (UC Davis sequencing facility). Plasmids containing variola virus DNA were handled and stored in the absence of replicating poxviruses.

### Luciferase-based reporter (LBR) assays

Assays were performed as described [9] with the following modifications: 5 × 10^4^ Hela PKR^kd^ cells per well 24 well plates were seeded 16 hours prior to the experiment. Cells were transfected with 200 ng of the indicated PKR expression vector, the indicated amount of each K3L expression vector, and 50 ng of pGL3 firefly luciferase expression vector (Promega) using GenJet (Signagen) at a DNA to GenJet ratio of 1:2 following the manufacturer’s protocol. Empty pSG5 vector was used to maintain the DNA concentration at 200 or 100 ng where appropriate. 48 hours post-transfection cells were lysed with mammalian lysis buffer (GE Healthcare), then luciferin (Promega) was added following the manufacturer’s recommendations. Luciferase activity was measured using a GloMax luminometer (Promega). Experiments were conducted in triplicate in three independent experiments.

### Viruses and infection assays

VACV Copenhagen strain VC2 and vP872 [13] were kindly provided by Dr. Bertram Jacobs. Generation of VC-R4, a derivative of vP872, was described [22]. VC-R4:VACV_K3L, VC-R4:CMLV_032 and VC-R4:CMLV_032A25V/V70I were generated by the scarless integration of the open reading frames of the K3L orthologs into the E3L locus in VC-R4 by the same method. The resulting viruses were plaque-purified three times and the integrations were confirmed by Sanger sequencing (UC Davis sequencing facility).

Plaque assays were performed with confluent six-well plates of the indicated cell lines, which were infected with 30 plaque forming units (pfu) of each indicated virus, as determined on RK13+E3L+K3L cells. One hour post-infection, the medium was replaced with DMEM containing 1% carboxymethylcellulose (CMC). After 48 hours, cells were washed with PBS and stained with 0.1% crystal violet. The plates were imaged using an iBright Imaging System (Invitrogen).

Multiple-cycle virus replication assays were carried out in confluent six-well plates of the indicated cells, which were infected with each indicated virus (MOI = 0.01). After 48 hours, cells and supernatants were collected and subjected to three rounds of freezing at -80 °C and thawing at 37 °C. Lysates were sonicated for 15s, 50% amplitude (Qsonica Q500). Viruses were titered by 10-fold serial dilutions on RK13+E3L+K3L cells. Infections and viral titer were performed in duplicate.

### Western blot analyses

To detect expression of K3 orthologs in transfected cells, 4 × 10^5^ Hela PKR-knock-down cells were seeded in six-well plates and transfected with 3 µg of the indicated K3-expressing plasmids following the GenJet protocol described above. After 48 h, cells were lysed with 1% sodium dodecyl sulfate (SDS) in PBS (VWR) and sonicated at 50% amplitude for 10 seconds twice. All protein lysates were separated on 12% SDS-polyacrylamide gels and transferred to polyvinyl difluoride (PVDF, GE Healthcare) membranes. Membranes were blocked with 5% non-fat milk dissolved in TBST (20M Tris, 150mM NaCl, 0.1% Tween 20, pH 7.4) for 1 hour. Blots were probed with a primary antibody (1:3000) directed against VACV K3 [17], or anti-β-actin (1:10,000, Sigma-Aldrich, A5441). All primary antibodies were diluted in TBST containing 5% BSA and 0.02% (w/v) sodium azide. Membranes were incubated overnight at 4°C in the primary antibody, washed with TBS-T three times for 5 mins, and then incubated for 1 hour at room temperature with donkey anti-rabbit or goat anti-mouse secondary antibodies conjugated to horseradish peroxidase (Invitrogen, A16110, 62–6520) at 1:10,000 in TBST containing 5% (w/v) nonfat milk. The membranes were then washed five times for 5 min and proteins were detected with Amersham ECL (GE Healthcare). Images were taken using the iBright Imaging System (Invitrogen).

### PKR phylogeny

The amino acid sequences of PKR from 40 different species were aligned using MUSCLE [47] and the ambiguously aligned positions were eliminated using Gblocks [48]. The phylogenetic analysis was performed using the maximum likelihood approach, as implemented in PhyML v3.0 [49]. Branch support was estimated using the aLRT statistics, and the phylogenetic tree was constructed using the default substitution model, as implemented in the automatic model selection in PhyML [50]. One hundred bootstrap replicates were performed to estimate the nodal support at major branch points. The resulting phylogenetic tree was visualized using FigTree.

## Acknowledgments

We thank Adam Geballe, Stephanie Child, Charles Samuel, Bertram Jacobs, Udeni Balasuriva, Ying Fang, and Nels Elde for cell lines, viruses and plasmids. This work was supported by grant AI114851 (to S.R.) from the National Institute of Allergy and Infectious Diseases, National Institutes of Health.

## Author contributions

C. Park, C. Peng and S.R. designed the experiments. C. Park, C. Peng, M.J.R., S.H., L.T. and G.B. performed experiments and analyzed data. C. Park, G.B., L.T. and S.R. wrote the manuscript. S.R. conceptualized and supervised the project.

## References

1. Haller SL, Peng C, McFadden G, Rothenburg S. Poxviruses and the evolution of host range and virulence. Infect Genet Evol. 2014;21:15–40. doi: 10.1016/j.meegid.2013.10.014. PubMed PMID: 24161410; PubMed Central PMCID: PMCPMC3945082.

2. Bera BC, Shanmugasundaram K, Barua S, Venkatesan G, Virmani N, Riyesh T, et al. Zoonotic cases of camelpox infection in India. Vet Microbiol. 2011;152(1-2):29-38. Epub 2011/05/17. doi: 10.1016/j.vetmic.2011.04.010. PubMed PMID: 21571451.

3. Khalafalla AI, Abdelazim F. Human and Dromedary Camel Infection with Camelpox Virus in Eastern Sudan. Vector Borne Zoonotic Dis. 2017;17(4):281-4. Epub 2017/01/06. doi: 10.1089/vbz.2016.2070. PubMed PMID: 28055328.

4. Pfaller CK, Li Z, George CX, Samuel CE. Protein kinase PKR and RNA adenosine deaminase ADAR1: new roles for old players as modulators of the interferon response. Curr Opin Immunol. 2011;23(5):573-82. Epub 2011/09/20. doi: 10.1016/j.coi.2011.08.009. PubMed PMID: 21924887; PubMed Central PMCID: PMCPMC3190076.

5. Wu S, Kaufman RJ. A model for the double-stranded RNA (dsRNA)-dependent dimerization and activation of the dsRNA-activated protein kinase PKR. J Biol Chem. 1997;272(2):1291-6. Epub 1997/01/10. PubMed PMID: 8995434.

6. Rothenburg S, Georgiadis MM, Wek RC. Evolution of eIF2α Kinases: Adapting Translational Control to Diverse Stresses. In: Hernández G, Jagus R, editors. Evolution of the Protein Synthesis Machinery and Its Regulation. Cham: Springer International Publishing; 2016. p. 235–60.

7. Langland JO, Cameron JM, Heck MC, Jancovich JK, Jacobs BL. Inhibition of PKR by RNA and DNA viruses. Virus Res. 2006;119(1):100-10. Epub 2006/05/18. doi: 10.3201/eid1204.051181. PubMed PMID: 16704884.

8. Elde NC, Child SJ, Geballe AP, Malik HS. Protein kinase R reveals an evolutionary model for defeating viral mimicry. Nature. 2009;457(7228):485-9. Epub 2008/12/02. doi: 10.1038/nature07529. PubMed PMID: 19043403; PubMed Central PMCID: PMCPMC2629804.

9. Rothenburg S, Seo EJ, Gibbs JS, Dever TE, Dittmar K. Rapid evolution of protein kinase PKR alters sensitivity to viral inhibitors. Nat Struct Mol Biol. 2009;16(1):63–70. doi: 10.1038/nsmb.1529. PubMed PMID: 19043413; PubMed Central PMCID: PMCPMC3142916.

10. Carpentier KS, Esparo NM, Child SJ, Geballe AP. A Single Amino Acid Dictates Protein Kinase R Susceptibility to Unrelated Viral Antagonists. PLoS Pathog. 2016;12(10):e1005966. Epub 2016/10/26. doi: 10.1371/journal.ppat.1005966. PubMed PMID: 27780231; PubMed Central PMCID: PMCPMC5079575.

11. Chang HW, Watson JC, Jacobs BL. The E3L gene of vaccinia virus encodes an inhibitor of the interferon-induced, double-stranded RNA-dependent protein kinase. Proc Natl Acad Sci U S A. 1992;89(11):4825-9. Epub 1992/06/01. PubMed PMID: 1350676; PubMed Central PMCID: PMCPMC49180.

12. Romano PR, Zhang F, Tan SL, Garcia-Barrio MT, Katze MG, Dever TE, et al. Inhibition of double-stranded RNA-dependent protein kinase PKR by vaccinia virus E3: role of complex formation and the E3 N-terminal domain. Mol Cell Biol. 1998;18(12):7304-16. Epub 1998/11/20. PubMed PMID: 9819417; PubMed Central PMCID: PMCPMC109312.

13. Beattie E, Tartaglia J, Paoletti E. Vaccinia virus-encoded eIF-2 alpha homolog abrogates the antiviral effect of interferon. Virology. 1991;183(1):419-22. Epub 1991/07/01. PubMed PMID: 1711259.

14. Dar AC, Sicheri F. X-ray crystal structure and functional analysis of vaccinia virus K3L reveals molecular determinants for PKR subversion and substrate recognition. Mol Cell. 2002;10(2):295-305. Epub 2002/08/23. PubMed PMID: 12191475.

15. Langland JO, Jacobs BL. The role of the PKR-inhibitory genes, E3L and K3L, in determining vaccinia virus host range. Virology. 2002;299(1):133-41. Epub 2002/08/09. PubMed PMID: 12167348.

16. Bratke KA, McLysaght A, Rothenburg S. A survey of host range genes in poxvirus genomes. Infect Genet Evol. 2013;14:406–25. doi: 10.1016/j.meegid.2012.12.002. PubMed PMID: 23268114; PubMed Central PMCID: PMCPMC4080715.

17. Park C, Peng C, Brennan G, Rothenburg S. Species-specific inhibition of antiviral protein kinase R by capripoxviruses and vaccinia virus. Ann N Y Acad Sci. 2019;1438(1):18-29. Epub 2019/01/16. doi: 10.1111/nyas.14000. PubMed PMID: 30644558.

18. Peng C, Haller SL, Rahman MM, McFadden G, Rothenburg S. Myxoma virus M156 is a specific inhibitor of rabbit PKR but contains a loss-of-function mutation in Australian virus isolates. Proc Natl Acad Sci U S A. 2016;113(14):3855–60. doi: 10.1073/pnas.1515613113. PubMed PMID: 26903626; PubMed Central PMCID: PMCPMC4833222.

19. Shchelkunov SN. An increasing danger of zoonotic orthopoxvirus infections. PLoS Pathog. 2013;9(12):e1003756. Epub 2013/12/18. doi: 10.1371/journal.ppat.1003756. PubMed PMID: 24339772; PubMed Central PMCID: PMCPMC3855571.

20. Mauldin MR, Antwerpen M, Emerson GL, Li Y, Zoeller G, Carroll DS, et al. Cowpox virus: What’s in a Name? Viruses. 2017;9(5). Epub 2017/05/10. doi: 10.3390/v9050101. PubMed PMID: 28486428; PubMed Central PMCID: PMCPMC5454414.

21. Franke A, Pfaff F, Jenckel M, Hoffmann B, Hoper D, Antwerpen M, et al. Classification of Cowpox Viruses into Several Distinct Clades and Identification of a Novel Lineage. Viruses. 2017;9(6). Epub 2017/06/13. doi: 10.3390/v9060142. PubMed PMID: 28604604; PubMed Central PMCID: PMCPMC5490819.

22. Vipat S, Brennan G, Park C, Haller SL, Rothenburg S. Rapid, Seamless Generation of Recombinant Poxviruses using Host Range and Visual Selection. J Vis Exp. 2020;(159). Epub 2020/06/09. doi: 10.3791/61049. PubMed PMID: 32510495.

23. Warren CJ, Sawyer SL. How host genetics dictates successful viral zoonosis. PLoS Biol. 2019;17(4):e3000217. Epub 2019/04/20. doi: 10.1371/journal.pbio.3000217. PubMed PMID: 31002666; PubMed Central PMCID: PMCPMC6474636.

24. Rothenburg S, Brennan G. Species-Specific Host-Virus Interactions: Implications for Viral Host Range and Virulence. Trends Microbiol. 2020;28(1):46-56. Epub 2019/10/11. doi: 10.1016/j.tim.2019.08.007. PubMed PMID: 31597598; PubMed Central PMCID: PMCPMC6925338.

25. Essbauer S, Pfeffer M, Meyer H. Zoonotic poxviruses. Vet Microbiol. 2010;140(3-4):229-36. Epub 2009/10/16. doi: 10.1016/j.vetmic.2009.08.026. PubMed PMID: 19828265.

26. Dabrowski PW, Radonic A, Kurth A, Nitsche A. Genome-wide comparison of cowpox viruses reveals a new clade related to Variola virus. PLoS One. 2013;8(12):e79953. Epub 2013/12/07. doi: 10.1371/journal.pone.0079953. PubMed PMID: 24312452; PubMed Central PMCID: PMCPMC3848979.

27. Antwerpen MH, Georgi E, Nikolic A, Zoeller G, Wohlsein P, Baumgartner W, et al. Use of Next Generation Sequencing to study two cowpox virus outbreaks. PeerJ. 2019;7:e6561. Epub 2019/03/09. doi: 10.7717/peerj.6561. PubMed PMID: 30847261; PubMed Central PMCID: PMCPMC6398431.

28. Lourie B, Nakano JH, Kemp GE, Setzer HW. Isolation of poxvirus from an African Rodent. J Infect Dis. 1975;132(6):677-81. Epub 1975/12/01. doi: 10.1093/infdis/132.6.677. PubMed PMID: 811713.

29. Arndt WD, White SD, Johnson BP, Huynh T, Liao J, Harrington H, et al. Monkeypox virus induces the synthesis of less dsRNA than vaccinia virus, and is more resistant to the anti-poxvirus drug, IBT, than vaccinia virus. Virology. 2016;497:125-35. Epub 2016/07/29. doi: 10.1016/j.virol.2016.07.016. PubMed PMID: 27467578; PubMed Central PMCID: PMCPMC5026613.

30. Frey TR, Lehmann MH, Ryan CM, Pizzorno MC, Sutter G, Hersperger AR. Ectromelia virus accumulates less double-stranded RNA compared to vaccinia virus in BS-C-1 cells. Virology. 2017;509:98-111. Epub 2017/06/20. doi: 10.1016/j.virol.2017.06.010. PubMed PMID: 28628829; PubMed Central PMCID: PMCPMC5541908.

31. Cao J, Varga J, Deschambault Y. Poxvirus encoded eIF2alpha homolog, K3 family proteins, is a key determinant of poxvirus host species specificity. Virology. 2020;541:101-12. Epub 2020/02/15. doi: 10.1016/j.virol.2019.12.008. PubMed PMID: 32056708.

32. Myskiw C, Arsenio J, Hammett C, van Bruggen R, Deschambault Y, Beausoleil N, et al. Comparative analysis of poxvirus orthologues of the vaccinia virus E3 protein: modulation of protein kinase R activity, cytokine responses, and virus pathogenicity. J Virol. 2011;85(23):12280-91. Epub 2011/09/16. doi: 10.1128/JVI.05505-11. PubMed PMID: 21917954; PubMed Central PMCID: PMCPMC3209343.

33. Rosengard AM, Liu Y, Nie Z, Jimenez R. Variola virus immune evasion design: expression of a highly efficient inhibitor of human complement. Proc Natl Acad Sci U S A. 2002;99(13):8808-13. Epub 2002/05/30. doi: 10.1073/pnas.112220499. PubMed PMID: 12034872; PubMed Central PMCID: PMCPMC124380.

34. Liszewski MK, Leung MK, Hauhart R, Buller RM, Bertram P, Wang X, et al. Structure and regulatory profile of the monkeypox inhibitor of complement: comparison to homologs in vaccinia and variola and evidence for dimer formation. J Immunol. 2006;176(6):3725-34. Epub 2006/03/07. doi: 10.4049/jimmunol.176.6.3725. PubMed PMID: 16517741.

35. Yadav VN, Pyaram K, Mullick J, Sahu A. Identification of hot spots in the variola virus complement inhibitor (SPICE) for human complement regulation. J Virol. 2008;82(7):3283-94. Epub 2008/01/25. doi: 10.1128/JVI.01935-07. PubMed PMID: 18216095; PubMed Central PMCID: PMCPMC2268448.

36. Yadav VN, Pyaram K, Ahmad M, Sahu A. Species selectivity in poxviral complement regulators is dictated by the charge reversal in the central complement control protein modules. J Immunol. 2012;189(3):1431-9. Epub 2012/06/27. doi: 10.4049/jimmunol.1200946. PubMed PMID: 22732591.

37. Kumar J, Yadav VN, Phulera S, Kamble A, Gautam AK, Panwar HS, et al. Species Specificity of Vaccinia Virus Complement Control Protein for the Bovine Classical Pathway Is Governed Primarily by Direct Interaction of Its Acidic Residues with Factor I. J Virol. 2017;91(19). Epub 2017/07/21. doi: 10.1128/JVI.00668-17. PubMed PMID: 28724763; PubMed Central PMCID: PMCPMC5599756.

38. Meng X, Chao J, Xiang Y. Identification from diverse mammalian poxviruses of host-range regulatory genes functioning equivalently to vaccinia virus C7L. Virology. 2008;372(2):372-83. Epub 2007/12/07. doi: 10.1016/j.virol.2007.10.023. PubMed PMID: 18054061; PubMed Central PMCID: PMCPMC2276162.

39. Liu J, Rothenburg S, McFadden G. The poxvirus C7L host range factor superfamily. Curr Opin Virol. 2012;2(6):764-72. Epub 2012/10/30. doi: 10.1016/j.coviro.2012.09.012. PubMed PMID: 23103013; PubMed Central PMCID: PMCPMC3508084.

40. Meng X, Krumm B, Li Y, Deng J, Xiang Y. Structural basis for antagonizing a host restriction factor by C7 family of poxvirus host-range proteins. Proc Natl Acad Sci U S A. 2015;112(48):14858-63. Epub 2015/11/19. doi: 10.1073/pnas.1515354112. PubMed PMID: 26578811; PubMed Central PMCID: PMCPMC4672790.

41. Liu J, Wennier S, Zhang L, McFadden G. M062 is a host range factor essential for myxoma virus pathogenesis and functions as an antagonist of host SAMD9 in human cells. J Virol. 2011;85(7):3270-82. Epub 2011/01/21. doi: 10.1128/JVI.02243-10. PubMed PMID: 21248034; PubMed Central PMCID: PMCPMC3067850.

42. Liu J, McFadden G. SAMD9 is an innate antiviral host factor with stress response properties that can be antagonized by poxviruses. J Virol. 2015;89(3):1925-31. Epub 2014/11/28. doi: 10.1128/JVI.02262-14. PubMed PMID: 25428864; PubMed Central PMCID: PMCPMC4300762.

43. Meng X, Zhang F, Yan B, Si C, Honda H, Nagamachi A, et al. A paralogous pair of mammalian host restriction factors form a critical host barrier against poxvirus infection. PLoS Pathog. 2018;14(2):e1006884. Epub 2018/02/16. doi: 10.1371/journal.ppat.1006884. PubMed PMID: 29447249; PubMed Central PMCID: PMCPMC5831749.

44. Zhang F, Meng X, Townsend MB, Satheshkumar PS, Xiang Y. Identification of CP77 as the Third Orthopoxvirus SAMD9 and SAMD9L Inhibitor with Unique Specificity for a Rodent SAMD9L. J Virol. 2019;93(12). Epub 2019/03/29. doi: 10.1128/JVI.00225-19. PubMed PMID: 30918078; PubMed Central PMCID: PMCPMC6613757.

45. Zhang P, Samuel CE. Protein kinase PKR plays a stimulus- and virus-dependent role in apoptotic death and virus multiplication in human cells. J Virol. 2007;81(15):8192-200. Epub 2007/05/25. doi: 10.1128/JVI.00426-07. PubMed PMID: 17522227; PubMed Central PMCID: PMCPMC1951329.

46. Rahman MM, Liu J, Chan WM, Rothenburg S, McFadden G. Myxoma virus protein M029 is a dual function immunomodulator that inhibits PKR and also conscripts RHA/DHX9 to promote expanded host tropism and viral replication. PLoS Pathog. 2013;9(7):e1003465. Epub 2013/07/16. doi: 10.1371/journal.ppat.1003465. PubMed PMID: 23853588; PubMed Central PMCID: PMCPMC3701710.

47. Edgar RC. MUSCLE: multiple sequence alignment with high accuracy and high throughput. Nucleic Acids Res. 2004;32(5):1792-7. Epub 2004/03/23. doi: 10.1093/nar/gkh340. PubMed PMID: 15034147; PubMed Central PMCID: PMCPMC390337.

48. Talavera G, Castresana J. Improvement of phylogenies after removing divergent and ambiguously aligned blocks from protein sequence alignments. Syst Biol. 2007;56(4):564-77. Epub 2007/07/27. doi: 10.1080/10635150701472164. PubMed PMID: 17654362.

49. Guindon S, Dufayard JF, Lefort V, Anisimova M, Hordijk W, Gascuel O. New algorithms and methods to estimate maximum-likelihood phylogenies: assessing the performance of PhyML 3.0. Syst Biol. 2010;59(3):307-21. Epub 2010/06/09. doi: 10.1093/sysbio/syq010. PubMed PMID: 20525638.

50. Lefort V, Longueville JE, Gascuel O. SMS: Smart Model Selection in PhyML. Mol Biol Evol. 2017;34(9):2422-4. Epub 2017/05/05. doi: 10.1093/molbev/msx149. PubMed PMID: 28472384; PubMed Central PMCID: PMCPMC5850602.

51. Dar AC, Dever TE, Sicheri F. Higher-order substrate recognition of eIF2alpha by the RNA-dependent protein kinase PKR. Cell. 2005;122(6):887-900. Epub 2005/09/24. doi: 10.1016/j.cell.2005.06.044. PubMed PMID: 16179258.

52. Kawagishi-Kobayashi M, Silverman JB, Ung TL, Dever TE. Regulation of the protein kinase PKR by the vaccinia virus pseudosubstrate inhibitor K3L is dependent on residues conserved between the K3L protein and the PKR substrate eIF2alpha. Mol Cell Biol. 1997;17(7):4146-58. Epub 1997/07/01. doi: 10.1128/mcb.17.7.4146. PubMed PMID: 9199350; PubMed Central PMCID: PMCPMC232268.

